# Structure-based design of hepatitis C virus E2 glycoprotein improves serum binding and cross-neutralization

**DOI:** 10.1101/2020.04.15.044073

**Authors:** Brian G. Pierce, Zhen-Yong Keck, Ruixue Wang, Patrick Lau, Kyle Garagusi, Khadija Elkholy, Eric A. Toth, Richard A. Urbanowicz, Johnathan D. Guest, Pragati Agnihotri, Melissa C. Kerzic, Alexander Marin, Alexander K. Andrianov, Jonathan K. Ball, Roy A. Mariuzza, Thomas R. Fuerst, Steven K.H. Foung

## Abstract

An effective vaccine for hepatitis C virus (HCV) is a major unmet need, and it requires an antigen that elicits immune responses to key conserved epitopes. Based on structures of antibodies targeting HCV envelope glycoprotein E2, we designed immunogens to modulate the structure and dynamics of E2 and favor induction of bNAbs in the context of a vaccine. These designs include a point mutation in a key conserved antigenic site to stabilize its conformation, as well as redesigns of an immunogenic region to add a new N-glycosylation site and mask it from antibody binding. Designs were experimentally characterized for binding to a panel of human monoclonal antibodies (HMAbs) and the coreceptor CD81 to confirm preservation of epitope structure and preferred antigenicity profile. Selected E2 designs were tested for immunogenicity in mice, with and without hypervariable region 1, which is an immunogenic region associated with viral escape. One of these designs showed improvement in polyclonal immune serum binding to HCV pseudoparticles and neutralization of isolates associated with antibody resistance. These results indicate that antigen optimization through structure-based design of the envelope glycoproteins is a promising route to an effective vaccine for HCV.

## Importance

Hepatitis C virus infects approximately 1% of the world’s population, and no vaccine is currently available. Due to the high variability of HCV and its ability to actively escape the immune response, a goal of HCV vaccine design is to induce neutralizing antibodies that target conserved epitopes. Here we performed structure-based design of several epitopes of the HCV E2 envelope glycoprotein to engineer its antigenic properties. Designs were tested in vitro and in vivo, demonstrating alteration of the E2 antigenic profile in several cases, and one design led to improvement of cross-neutralization of heterologous viruses. This represents a proof of concept that rational engineering of HCV envelope glycoproteins can be used to modulate E2 antigenicity and optimize a vaccine for this challenging viral target.

## Introduction

Hepatitis C virus (HCV) infection is a major global disease burden, with 71 million individuals, or approximately 1% of the global population, chronically infected worldwide, and 1.75 million new infections per year (1). Chronic HCV infection can lead to cirrhosis and hepatocellular carcinoma, the leading cause of liver cancer, and in the United States HCV was found to surpass HIV and 59 other infectious conditions as a cause of death (2). While the development of direct-acting antivirals has improved treatment options considerably, several factors impede the effective use of antiviral treatment such as the high cost of antivirals, viral resistance, occurrence of reinfections after treatment cessation, and lack of awareness of infection in many individuals since HCV infection is considered a silent epidemic. Therefore, development of an effective preventative vaccine for HCV is necessary to reduce the burden of infection and transmission, and for global elimination of HCV (3).

Despite decades of research resulting in several HCV vaccine candidates tested in vivo and in clinical trials (4, 5), no approved HCV vaccine is available. There are a number of barriers to the development of an effective HCV vaccine, including the high mutation rate of the virus which leads to viral quasi-species in individuals and permits active evasion of T cell and B cell responses (6). Escape from the antibody response by HCV includes mutations in the envelope glycoproteins, as observed in vivo in humanized mice (7), studies in chimpanzee models (8), and through analysis of viral isolates from human chronic infection (9). This was also clearly demonstrated during clinical trials of a monoclonal antibody, HCV1, which in spite of its targeting a conserved epitope on the viral envelope, failed to eliminate the virus, as viral variants with epitope mutations emerged under immune pressure and dominated the rebounding viral populations in all treated individuals (10, 11).

There have been a number of successful structure-based vaccine designs for variable viruses such as influenza (12, 13), HIV (14, 15), and RSV (16, 17) where rationally designed immunogens optimize presentation of key conserved epitopes, mask sites using N-glycans, or stabilize conformations or assembly of the envelope glycoproteins. Recent studies have reported use of several of these strategies in the context of HCV glycoproteins, including removal or modification of N-glycans to improve epitope accessibility (18, 19), removal of hypervariable regions (18, 20, 21), or presentation of key conserved epitopes on scaffolds (22, 23). However, such studies have been relatively limited compared with other viruses, in terms of design strategies employed and number of designs tested, and immunogenicity studies have not shown convincing improvement of glycoprotein designs over native glycoproteins in terms of neutralization potency or breadth (18, 21), with the possible exception of an HVR-deleted high molecular weight form of the E2 glycoprotein that was tested in guinea pigs (20).

Here we report the generation, characterization, and in vivo immunogenicity of novel structure-based designs of the HCV E2 glycoprotein, which is the primary target of the antibody response to HCV and a major vaccine target. Designs were focused on antigenic domain D, which is a key region of E2 targeted by broadly neutralizing antibodies (bNAbs) that are resistant to viral escape (24), as well as antigenic domain A, which is targeted by non-neutralizing antibodies (25, 26). Based on the intrinsic flexibility of the neutralizing face of E2 (27), which includes antigenic domain D, and on the locations of bNAb epitopes to this domain (24), we identified a structure-based design substitution to reduce the mobility of that region and preferentially form the bnAb-bound conformation. We also tested several substitutions to hyperglycosylate and mask antigenic domain A located in a unique region on the back layer of E2, as determined by fine epitope mapping (28), which represents an approach that has been applied to mask epitopes in influenza (29) and HIV (30) glycoproteins. Designs were tested for antigenicity using a panel of monoclonal antibodies (mAbs), and selected designs were tested individually and as combinations for in vivo immunogenicity. Assessment of immunized sera revealed that certain E2 designs yielded improvements in serum binding to recombinant HCV particles, as well as viral cross-neutralization, while maintaining serum binding to soluble E2 glycoprotein and key epitopes. This provides a proof-of-concept that rational design of HCV glycoproteins can lead to improvements in immunogenicity and neutralization breadth.

## Results

### Structure-based design of E2

We utilized two approaches to design variants of the E2 glycoprotein to improve its antigenicity and immunogenicity (Figure 1). For one approach, we used the previously reported structure of the affinity matured bnAb HC84.26.5D bound to its epitope from E2 antigenic domain D (31) (PDB code 4Z0X), which shows the same epitope conformation observed in the context of other domain D human monoclonal antibodies (HMAbs) targeting this site (32). Analysis of this epitope structure for potential proline residue substitutions to stabilize its HMAb-bound conformation identified several candidate sites (Figure 1A, Table 1). We selected one of these substitutions, H445P, that is adjacent to core contact residues for domain D located at aa 442-443 (32) for subsequent experimental characterization, due to its position in a region with no secondary structure, and location between residues Y443 and K446 which both make key antibody contacts in domain D antibody complex structures (31, 32). This also represents a distinct region of the epitope from a substitution that we previously described and tested (A439P) (28).

**Figure 1.**
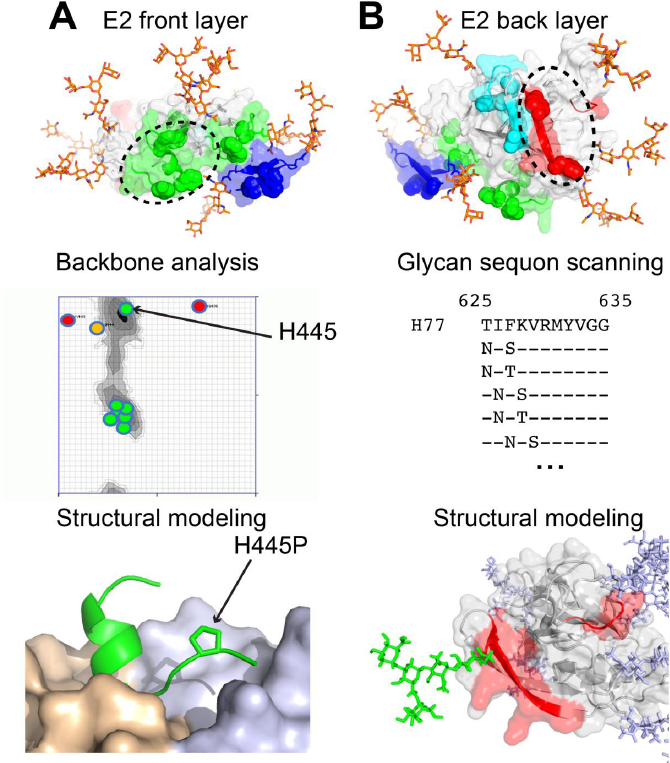
Structure-based design of E2 to stabilize and mask epitopes. A) Design of the E2 front layer. (top) Antigenic domain B-D supersite (green; also referred to as antigenic region 3) is highlighted on the E2 core X-ray structure (63), along with antigenic domain E (blue), and modeled N-glycans shown as orange sticks. Ramachandran plot analysis for proline-like backbone conformation (middle) and structural modeling of proline substitution structural and energetic effects (bottom) were performed using RosettaDesign and the HC84.26.5D-AS434 epitope complex structure (31). HC84.26.5D HMAb is shown as wheat and light blue surface, epitope is green with selected mutant residue (H445P) shown as sticks. B) Design of the E2 back layer. (top) Antigenic domain A (red) was targeted for design and is shown on the E2 core structure, and antigenic domain C and modeled N-glycans are shown in cyan and orange sticks, respectively. (middle) Computational N-glycan scanning of antigenic domain A residues was performed to identify substitutions to mask its surface with designed NxS and NxT sequons. (bottom) Modeling of sequon mutants was performed in Rosetta (64), followed by modeling of N-glycan structures in the Glyprot Server (55). Modeled N-glycan design at Y632 (Y632N-G634S) shown as green sticks, with E2 core structure gray (based on H77C E2, PDB code 4MWF), E2 antigenic domain A residues red, and modeled native E2 N-glycans light blue sticks.

**Table 1.** Backbone structure and proline mutant analysis of antigenic domain D residues.

| Residue | Wild-type aa | Backbone angles<sup>1</sup> |  | Backbone analysis<sup>1</sup> |  | Rosetta |
| --- | --- | --- | --- | --- | --- | --- |
| | | $\Phi, ^\circ$ | $\Psi, ^\circ$ | Pro | Pre-Pro | Proline $\Delta\Delta G^2$ |
| 437 | Trp | -73.66 | -18.28 | Yes | No | 0.5 |
| 438 | Leu | -59.18 | -59.67 | Yes | No | 0.3 |
| 439 | Ala | -59.14 | -37.12 | Yes | Yes | 0 |
| 440 | Gly | -56.62 | -26.31 | Yes | Yes | 0.1 |
| 441 | Leu | -68.3 | -40.87 | Yes | Yes | 2.1 |
| 442 | Phe | -80.55 | -40.18 | Yes | Yes | 2.7 |
| 443 | Tyr | -159.11 | 143.78 | No | Yes | 2.5 |
| 444 | Gln | -110.46 | 128.63 | No | Yes | 2.2 |
| 445 | His | -58.15 | 153.22 | Yes | Yes | 0 |

<sup>1</sup>Values and proline backbone analysis were obtained from the Ramachandran plot analysis web server (http://zlab.bu.edu/rama)(53). Pre-Pro assessments correspond to pre-proline Ramachandran plot conformation for the backbone of the preceding residue. Unfavorable Pro or Pre-Pro conformations are noted with gray shaded cells.
<sup>2</sup>Predicted binding energy change for proline epitope mutant to the HC84.26.5D HMAb, based on the X-ray structure of the complex (PDB code 4Z0X) and computational mutagenesis in Rosetta (54). Predicted destabilizing values ( $> 0.5$ in Rosetta energy units) are indicated with shaded cells.

Another design approach, hyperglycosylation, was utilized to mask antigenic domain A, which is an immunogenic region on the back layer of E2 associated with non-neutralizing antibodies (25, 26, 28). Other antibodies with some binding determinants mapped to this region, including HMAbs AR1A and HEPC46, exhibit limited or weak neutralization (33). NxS (Asparagine-X-Serine) and NxT (Asparagine-X-Threonine) N-glycan sequon substitutions were modeled in Rosetta at solvent-exposed E2 positions in antigenic domain A (Figure 1B, Table 2), followed by visual inspection of the modeled E2 mutant structures to confirm exposure of the mutant asparagine residues. This analysis suggested that designs with N-glycans at residues 627 (F627N-V629T), 628 (K628N-R630S), 630 (R630N-Y632T), and 632 (Y632N-G634S) warranted further investigation for effects on antigenicity.

**Table 2.** Calculated surface accessibility of E2 residues in antigenic domain A.

| **Residue** | **Amino Acid** | **Side Chain ASA<sup>1</sup>** |
| --- | --- | --- |
| 627 | Phe | 62.7 |
| 628 | Lys | 89.4 |
| 629 | Val | 38.4 |
| 630 | Arg | 60.4 |
| 631 | Met | 14 |
| 632 | Tyr | 62.3 |
| 633 | Val | 1.5 |

<sup>1</sup>Accessible surface areas calculated using the X-ray structure of H77 E2 core (PDB code 4MWF) and the naccess program (56) with default parameters. Values reflect relative side chain surface accessibility (normalized to 100).

### Characterization of mutant antigenicity using ELISA

We first screened the structure-based designs described above to assess their effects on E2 glycoprotein antigenicity, to confirm that designs preserved the structure of key E2 epitopes, and to disrupt non-neutralizing antigenic domain A HMAb binding in the case of the N-glycan designs. These designs were cloned in E1E2 and assessed using ELISA with a panel of representative HMAbs to antigenic domains A-E (Figure 2). The results indicated that mutant H445P maintained approximately wild-type levels of binding to antibodies, while truncations of HVR1 had varying effects. Binding of domain E HMAb HC33.4, and to a lesser extent HC33.1, was negatively affected by truncation of all of HVR1 (residues 384-410 removed; referred to here as ΔHVR1_411_), whereas a more limited HVR1 truncation (residues 384-407 removed; referred to here as ΔHVR1) largely restored binding of these bNAbs. The design of ΔHVR1 was based on the observation that residue 408 located within HVR1 affected the binding of HC33.4 but not HC33.1 (34). Likewise, designed N-glycan substitutions showed varying effects on antigenicity, with pronounced reduction of binding for several bNAbs for F627NT (F627N-V629T) and R630NT (R630N-Y632T), while K628NS (K628N-R630S) did not exhibit ablation of domain A antibody binding. In contrast, Y632NS (Y632N-G634S) disrupted binding for both tested domain A HMAbs, with limited loss of binding for other HMAbs. Based on this antigenic characterization, designs H445P, ΔHVR1, and Y632NS were selected for further testing.

**Figure 2.**
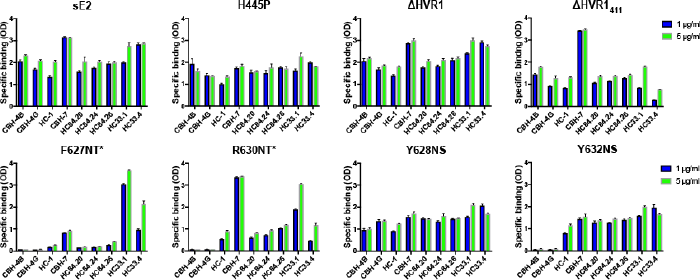
Antigenic characterization of E2 designs using ELISA. Designs were cloned and expressed in the context of E1E2 as previously described (28) and tested in for binding to a panel of HMAbs that target E2 antigenic domain A (CBH-4G, CBH-4B), B (HC-1), C (CBH-7), D (HC84.28, HC84.24, HC84.26), and E (HC33.1, HC33.4), at concentrations of 1 μg/ml, and 5 μg/ml. Binding was tested to wild-type H77 E2 (sE2; residues 384-661), and compared with sE2 designs ΔHVR1 (residues 408-661), ΔHVR1_411_ (residues 411-661), H445P, V627NT (V627N-V629T), R630NT (R630N-Y632T), K628NS (K628N-R630S), and Y632NS (Y632N-G634S). Asterisks denote designs that were tested in the context of ΔHVR1_411_ (aa 411-661) sE2 rather than full length sE2 (384-661).

### Biophysical and antigenic characterization of E2 designs

The two candidate structure-based E2 designs H445P, Y632NS, as well as ΔHVR1, were expressed as soluble E2 (sE2) glycoproteins and tested for thermostability and binding affinity to a panel of HMAbs, as well as the CD81 receptor (Table 3). Pairwise combinations of these designs, and a “Triple” design with all three modifications, were also expressed and tested. As noted previously by others (27), wild-type sE2 was found to exhibit high thermostability (T_m_ = 84.5 °C in Table 3). All designs likewise showed high thermostability, with only minor reductions in T_m_, with the exception of combined Triple which had the lowest measured thermostability among the tested E2 mutants (T_m_ = 76.5 °C).

**Table 3.** Antigenic and biophysical characterization of E2 designs.

| Antibody/Receptor Binding K<sub>D</sub>, nM<sup>3</sup> |  |  |  |  |  |  |  |  |  |  |
| --- | --- | --- | --- | --- | --- | --- | --- | --- | --- | --- |
| sE2 Construct<sup>1</sup> | T<sub>m</sub><sup>2</sup> | Domain A |  | Domain B |  | Domain D |  | Domain E |  | Receptor |
|  |  | CBH-4D | CBH-4G | AR3A | HEPC 74 | HC84.26.WH.5DL | HC84.1 | HCV-1 | HC33.1 | CD81 |
| sE2 wild-type | 84.5 | 8.4 | 15 | 1.8 | 5.1 | 4.3 | 30 | 42 | 9.5 | 22 |
| H445P | 83.1 | 14 | 15 | 1.1 | 4.6 | **0.4** | 31 | 70 | 8.0 | 23 |
| Y632NS | 81.4 | 37 | **180** | 1.2 | 4.6 | 4.0 | 83 | 26 | 6.5 | 31 |
| ΔHVR1 | 84.5 | 5.8 | 20 | 1.5 | 7.0 | 3.1 | 60 | 42 | 7.9 | 15 |
| ΔHVR1-H445P | 82.9 | 11 | 15 | 1.5 | 2.3 | 2.4 | 51 | 46 | 7.6 | 10 |
| ΔHVR1-Y632NS | 80.0 | 7.7 | **140** | 1.8 | 9.0 | 2.2 | 110 | 50 | 6.7 | **140** |
| Triple | 76.5 | 22 | **110** | 1.4 | 4.6 | 2.8 | 58 | 27 | 9 | 40 |

<sup>1</sup>sE2 wild-type corresponds to residues 384-661 of H77C E2, and listed designs represent point mutants or truncations of that sequence. Y632NS is an abbreviation for the double mutant Y632N-G634S, and ΔHVR1 denotes deletion of most of the HVR1 sequence at the N-terminus of E2, with resultant construct containing residues 408-661. “Triple” denotes combination of ΔHVR1, H445P, and Y632NS designs.
<sup>2</sup>T<sub>m</sub> values are in °C and were measured by differential scanning calorimetry.
<sup>3</sup>Steady-state dissociation constant (K<sub>D</sub>) values were measured by Octet biolayer interferometry. Antibodies are classified by their mapping to antigenic domains A, B, D, and E on E2 (28). Values in bold denote K<sub>D</sub> changes more than 5-fold versus wild-type E2 for that antibody.

To assess antigenicity of glycoprotein designs, solution binding affinity measurements were performed with Octet using HMAbs that target E2 antigenic domains A, B, D, and E, with two antibodies per domain, as well as the receptor CD81 (Table 3). These antibodies have been previously characterized using multiple global alanine scanning studies (28, 35) (CBH-4G, CBH-4D, HC33.1, AR3A, HC33.1), and X-ray structural characterization studies (AR3A, HEPC74, HC84.1, HC33.1, HCV1) (32, 36-39). The HC84.26.WH.5DL is an affinity matured clone of the parental HC84.26 antibody with improved affinity and neutralization breadth over the parental antibody (31). The binding site of CD81 has been mapped to E2 residues in antigenic domains B, D, and E (35), thus CD81 binding provides additional assessment of antigenicity of that E2 supersite (6). Binding experiments with this panel showed nanomolar binding affinities to wild-type sE2, which were largely maintained for sE2 designs. A 10-fold increase in binding affinity of sE2 design H445P for domain D HMAb HC84.26.WH.5DL was observed, showing that this design, located within antigenic domain D, not only maintained affinity, but improved engagement in that case; a steady-state binding fit for that interaction is shown in Figure 3A. However, this effect was not observed for combinations of designs including H445P, suggesting possible interplay between designed sites. As expected, domain A hyperglycosylation designs Y632NS, ΔHVR1-Y632NS, and Triple (ΔHVR1-H445P-Y632NS) showed loss of binding (>5-fold for each) to antigenic domain A HMAb CBH-4G (Y632NS-CBH-4G binding measurement is shown in Figure 3B), though we did not observe disruption of binding to CBH-4D. Additionally, design ΔHVR1-Y632NS showed moderate (6-fold) loss of CD81 binding, which was not the case for other designs. As domain A HMAbs have distinct, albeit similar, binding determinants on E2 (28), differential effects on domain A antibody binding by Y632NS variants reflect likely differences in HMAb docking footprints on E2. Measurements of glycan occupancy at residue 632 using mass spectroscopy showed partial levels of glycosylation at that site for Y632NS and combinations (Table 4), which may be responsible for incomplete binding ablation to the tested antigenic domain A HMAbs. As alanine substitution at Y632 was previously found to disrupt binding of domain A antibodies (28), it is possible that the Y632N amino acid substitution in the Y632NS may be responsible, in addition to partial N-glycosylation, for effects on domain A antibody binding. Regardless, these results suggest at least partial binding disruption and N-glycan masking of this region, supporting testing of those designs as immunogens in vivo.

**Figure 3.**
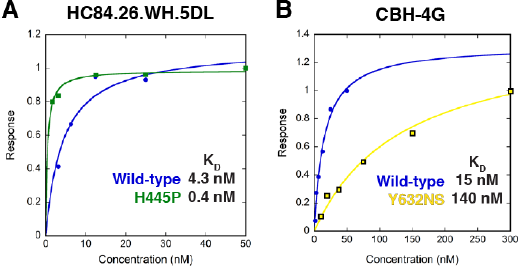
Antigenic characterization of sE2 designs H445P and Y632NS using biolayer interferometry (BLI). (A) Measured binding of broadly neutralizing monoclonal antibody HC84.26.WH.5DL to E2 design H445P compared to wild-type soluble E2 (sE2). (B) Measured binding of non-neutralizing monoclonal antibody CBH-4G to E2 design Y632NS (Y632N-G634S) compared to wild-type soluble E2 (sE2). Steady-state binding curve fits are shown, which were used to determine binding dissociation constants (K_d_) values.

**Table 4.** Percentage occupancy for engineered N-glycan at position 632, determined by mass spectrometry.

| **Design** | **% Glycosylation** | **EndoF Efficiency** |
| --- | --- | --- |
| Y632NS | 51 | 96 |
| ΔHVR1-Y632NS | 44 | 97 |
| Triple<sup>1</sup> | 22 | 100 |

<sup>1</sup>Combination of ΔHVR1, H445P, and Y632NS.

### In vivo immunogenicity of E2 designs

Following confirmation of antigenicity, E2 designs were tested in vivo for immunogenicity, to assess elicitation of antibodies that demonstrate potency and neutralization breadth. CD1 mice (6 per group) were immunized with H77C sE2 and designs, using with Day 0 prime followed by three biweekly boosts. Sera were obtained at Day 56 (two weeks after the final boost) and tested for binding to H77C sE2 and key conserved epitopes (AS412/Domain E, AS434/Domain D) (Figure 4). Peptide epitopes were confirmed for expected monoclonal antibody specificity using ELISA (Figure 4B). Endpoint titers demonstrated that sera from mice immunized with E2 designs maintained recognition of sE2 and tested epitopes. Intra-group variability resulted in lack of statistically significant differences in serum binding between immunized groups, however mean titers from ΔΗVR1 group were moderately lower than the wild-type sE2 group, and other mutants yielded moderately higher serum binding to the tested epitopes. Notably, design H445P elicited antibodies that robustly cross-reacted with the wild type AS434/Domain D epitope. To assess differential binding to conformational epitopes on E2, serum binding competition with selected HMAbs was performed (Figure S1) but did not show major differences between immunized groups.

**Figure 4.**
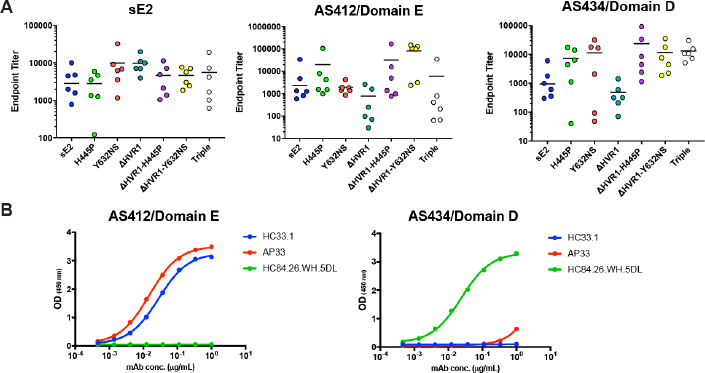
Immunized serum recognition of E2 and two E2 epitopes. A) Immunized sera were tested using ELISA for binding to soluble H77C E2 (sE2) and linear epitopes from antigenic domain E (AS412, aa 410-425) and antigenic domain D (AS434, aa 434-446). Serum binding was tested at successive three-fold dilutions starting at 1:60, and values are reported as endpoint titers. B) Binding of peptides to control monoclonal antibodies HC33.1 (58), AP33 (65), and HC84.26.WH.5DL (31).

### Serum binding to HCV E1E2 and HCV pseudoparticles

For further analysis of immunized serum binding, we tested binding to purified recombinant H77C E1E2 and HCV pseudoparticles from H77C and two heterologous genotypes (Figure 5, Table 5). While binding to H77 E1E2 resembled binding to H77 sE2, with no apparent difference between immunized groups, we observed notable differences in binding to HCVpps representing H77C, UKNP1.18.1, and J6 for H445P-immunized mice versus mice immunized with wild-type sE2. The difference between J6 HCVpp binding from H445P-immunized mice versus sE2-immunized mice was highly significant (p ≤ 0.0001, Kruskal-Wallis test). To confirm this difference in HCVpp binding between sE2 and H445P immunized groups, given the relatively low levels of overall titers, H77C HCVpps were purified and tested in ELISA for binding to pooled sera from sE2 and H445P immunized mice. This confirmed differences between immunized groups for sera from Day 56, as well as Day 42, which corresponds to three rather than four immunizations (Figure 6). To demonstrate native-like E2 and E1E2 assembly of the HCVpps in the context of the ELISA assay, purified HCVpps showed binding to monoclonal antibodies that target linear and conformational epitopes on E2 (HCV1, HC84.26.WH.5DL, AR3A) and conformational epitopes on E1E2 (AR4A, AR5A), and did not interact with negative control antibody (CA45) (Figure S2). The molecular basis for the differential serum reactivity when using HCVpp versus purified recombinant E1E2 in ELISA is unclear, particularly given that sE2 was used as an immunogen, yet these results collectively provide evidence that H445P may improve targeting of conserved glycoprotein epitopes on the intact HCV virion.

**Figure 5.**
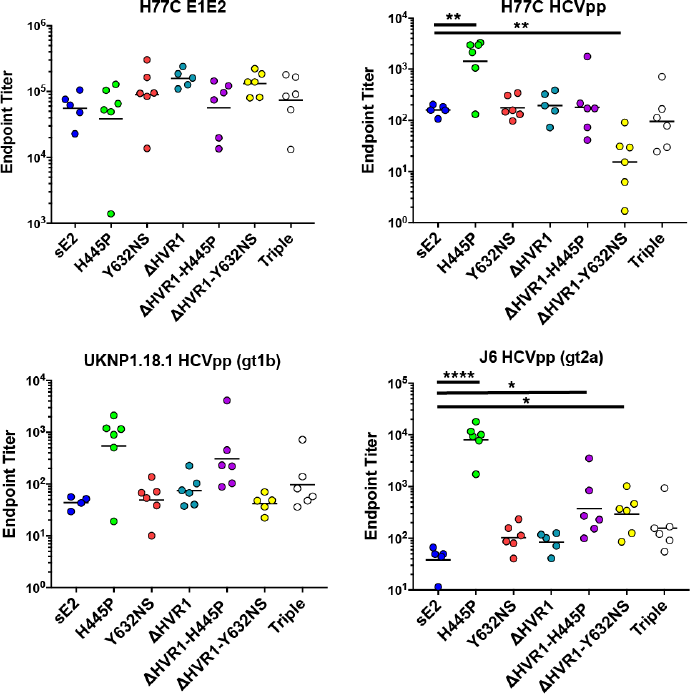
Immunized serum binding to recombinant E1E2 and HCV pseudoparticles (HCVpp). Immunized sera were tested for binding to H77C E1E2 and HCVpp representing H77C, UKNP1.18.1, and J6 isolates using ELISA. Serum binding was tested at three-fold dilutions starting at 1:100, and values are reported as endpoint titers. Due to insufficient sera, endpoint titers are not available for one mouse in the sE2 group and one mouse in ΔHVR1 group (E1E2, H77C HCVpp, J6 HCVpp), as well as two mice in the sE2 group and one mouse in the ΔHVR1-Y632NS group (UKNP1.18.1 HCVpp). P-values between group endpoint titer values were calculated using Kruskal-Wallis analysis of variance with Dunn’s multiple comparison test, and significant p-values between sE2 control and sE2 design groups are shown (*: p ≤ 0.05; **: p ≤ 0.01; ****: p ≤ 0.0001).

**Table 5.** Panel of viral isolates used in neutralization assays.

| Isolate | Genotype | Neutralization Phenotype | Neutralization Resistance Rank | % ID H77 sE2 | % ID H77 ΔHVR1 |
| --- | --- | --- | --- | --- | --- |
| H77 | 1a | Moderate | 22 | 100 | 100 |
| UKNP1.18.1 | 1b | Resistant | 3 | 79 | 82 |
| UKNP1.20.3 | 1b | Moderate | 32 | 79 | 82 |
| UKNP2.4.1 | 2a | Resistant | 1 | 72 | 74 |
| UKNP2.1.2 | 2i | Moderate | 23 | 70 | 73 |
| UKNP4.1.1 | 4a | Resistant | 2 | 71 | 75 |
| J6 | 2a | Resistant | 11 | 71 | 74 |

<sup>1</sup>Neutralization phenotype and neutralization resistance rank based on assessment of 78 HCVpp with a panel of monoclonal antibodies by Urbanowicz et al. (66).
<sup>2</sup>Percent identities reflect percent amino acid sequence identities with H77C sE2 (aa 384-661) and ΔHVR1 (aa 408-661).

**Figure 6.**
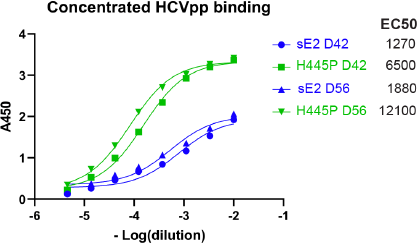
Comparison of purified HCV pseudoparticle (HCVpp) binding of immunized mouse sera from sE2 wild-type and H445P immunization. Purified H77C HCVpp were tested for binding to pooled murine sera from sE2 wild-type and H445P groups using ELISA, for sera from Day 42 and Day 56. Best-fit curves are shown and were used to calculate EC50 values.

### Homologous and heterologous serum neutralization

To assess effects of antibody neutralization potency and breadth from E2 designs, we tested serum neutralization of HCVpp representing homologous H77C (Figure 7) and six heterologous isolates (Figure 8). The heterologous isolates collectively diverge substantially in sequence from H77C and represent neutralization phenotypes ranging from moderately to highly resistant (Table 5). As we found previously, there was relatively large intra-group variability in neutralization of H77C (19), and no statistically significant differences between groups were observed. However, heterologous isolates showed much less intra-group variability, with some cases of designs yielding significantly higher neutralization than wild-type sE2. Notably, two resistant isolates had significantly higher neutralization for H445P-immunized sera than wild-type sE2-immunized sera (UKNP1.18.1, J6).

**Figure 7.**
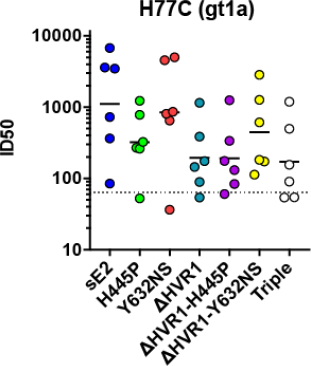
Serum neutralization of homologous (H77C) HCVpp. Neutralization titers are represented as serum dilution levels required to reach 50% virus neutralization (ID50), calculated by curve fitting in Graphpad Prism software. Serum dilutions were performed as two-fold dilutions starting at 1:64, and minimum dilution level (corresponding to 1:64) is indicated by a dotted line for reference.

**Figure 8.**
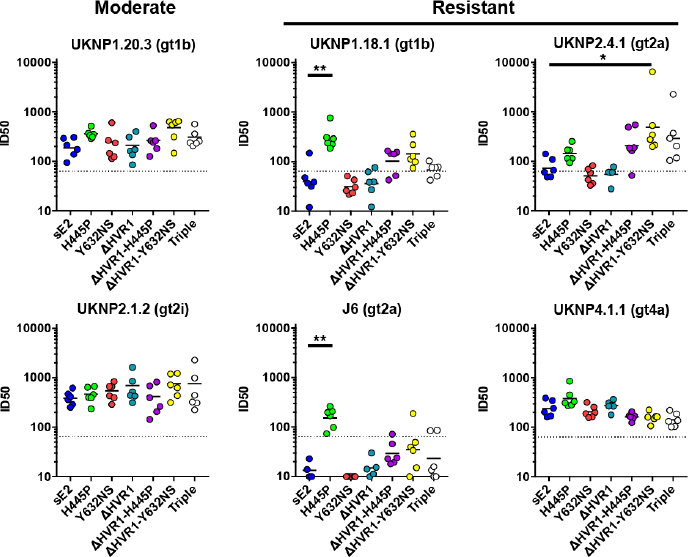
Serum neutralization of heterologous HCV HCVpp. Immunized murine serum neutralization was tested using HCV pseudoparticles (HCVpps) representing six heterologous isolates. Neutralization for four HCVpp representing isolates with resistant phenotypes are shown on the right, as indicated. Neutralization titers are represented as serum dilution levels required to reach 50% virus neutralization (ID50), calculated by curve fitting in Graphpad Prism software. Serum dilutions were performed as two-fold dilutions starting at 1:64, and minimum dilution levels (corresponding to 1:64) are indicated as dotted lines for reference. Murine sera with low (ID50 < 10) or incalculable ID50 values due to low or background levels of neutralization (observed only for some mice for J6 HCVpp neutralization) have ID50 shown as 10. Due to insufficient sera, J6 neutralization measurements did not include two mice from group 1 (sE2) and one mouse from group 4 (ΔHVR1). P-values between group ID50 values were calculated using Kruskal-Wallis analysis of variance with Dunn’s multiple comparison test, and significant p-values between sE2 control and sE2 design groups are shown (*: p ≤ 0.05; **: p ≤ 0.01).

## Discussion

In this study, we applied a variety of rational design approaches to engineer the HCV E2 glycoprotein to improve its antigenicity and immunogenicity. One of these approaches, removal of HVR1 (ΔHVR1), has been tested in several recent immunogenicity studies, in the context of E2 (18, 20, 23) and E1E2 (21). In this study, we tested the E2 ΔHVR1 mutant with residues 384-407 removed, which retains residues 408-661 of E2; this is a more conservative truncation than previously tested ΔHVR1 mutants, in order to retain residue 408 which is binding determinant for the HC33.4 HMAb and others (28, 34). Here we found this mutant to not be advantageous from an immunogenicity standpoint, which is in agreement with most other previous immunogenicity studies testing ΔHVR1 mutants (18, 21, 23). Although HVR1 is an immunogenic epitope, its removal from recombinant E2 glycoprotein does not appear to increase homologous or heterologous nAb titers, with the latter suggesting that the level of antibodies targeting conserved nAb epitopes did not increase upon HVR1 removal. Based largely on studies of engineered viruses in cell culture, as summarized in a recent review (40), removal of HVR1 is associated with increased nAb sensitivity and CD81 receptor binding, while a recent study has indicated that HVR1 may modulate viral dynamics and open and closed conformations during envelope breathing (41). Despite its importance in the context of the virion and its dynamics, its removal appears to have a neutral or minimal effect on the immunogenicity of recombinant envelope glycoproteins.

Another design strategy tested in this study was hyperglycosylation, through structure-based addition of N-glycan sequons to mask antigenic domain A, which is associated with non-neutralizing antibodies (25, 26, 28, 42). The concept of down-modulating immunity to this region was based on the observation that this region is highly immunogenic and may divert antibody responses to bNAb epitopes of lower immunogenicity. Through the efforts of isolating bNAbs to distinct regions on E2 from multiple HCV infected individuals, non-neutralizing antibodies to domain A are consistently identified (personal communication, S. Foung). This strategy has been successfully employed for other glycoprotein immunogens, including for HIV Env SOSIP trimers, where the immunogenic V3 loop was masked with designed N-glycans (30). Surprisingly, some of the designs in this study exhibited an impact on recognition by antibodies targeting antigenic domain D on the front layer of E2, suggesting a possible interplay between the front and back layers of E2, as proposed previously based on global alanine scanning mutagenesis (35). As observed by Ringe et al. in the context of HIV Env (30), the designed E2 N-glycan variant tested for immunogencity in this study (Y632NS) did not show improvements in nAb elicitation. However, its combination with ΔHVR1 did lead to modest improvement in nAb titers against one resistant isolate (UKNP2.4.1; p-value < 0.05), compared with wild-type sE2. Previously we used insect cell expression to alter the N-glycan profile of sE2 versus mammalian cell expressed sE2 (19), and others have recently tested immunogenicity for glycan-deleted E2 and E1E2 variants (18); in neither case was a significant improvement in homologous and heterologous nAb responses observed for immunogens with altered glycans. Collectively, these results suggest that glycoengineering of E2 or E1E2 represents a more challenging, and possibly less beneficial, avenue for HCV immunogen design, however a report of success by others through insect cell expressed sE2 indicates that altered glycosylation may help in some instances (43).

The designed substitution H445P, which was generated to preferentially adopt the bnAb-bound form in a portion of E2 antigenic domain D that exhibits structural variability (31), showed the greatest level of success, both with regard to improvements in serum binding to homologous and heterologous HCVpp, as well as HCVpp neutralization of heterologous HCVpp. This design lies within a supersite of E2 associated with many broadly neutralizing antibodies (5, 6, 44, 45), and through biophysical characterization and molecular dynamics simulation experiments, others have found that this region is likely quite flexible (27, 46), providing a rationale for stabilizing key residues to engage and elicit bNAbs. Interestingly, a residue adjacent to the site of this design appears to be functionally important, with the Q444R substitution restoring viral infectivity in the context of an HCVpp with a domain E “glycan shift” substitution, N417S (8). The design strategy of utilizing proline residue substitutions to stabilize conformations of viral glycoproteins has been successful for HIV Env (47), respiratory syncytial virus (RSV) F (48), MERS coronavirus spike (49), and recently, the novel coronavirus (SARS-CoV-2) spike (50). The data from this study suggest that this approach is also useful in the context of HCV E2, and possibly E1E2.

This study provides a proof-of-concept for computational structure-based design of the HCV E2 glycoprotein to modulate its antigenicity and immunogenicity. Future studies with the H445P design include testing of its antigenicity and immunogenicity in the context of HCV E1E2, testing immunogenicity in other animal models, as well as confirmation of its impact on E2 structure through high resolution X-ray structural characterization and additional biophysical characterization. Furthermore, additional designed proline substitutions in this flexible E2 “neutralizing face” supersite may confer greater improvements in homologous and heterologous nAb elicitation; these can be generated using structure-based design, or with a semi-rational library-based approach, as was used to scan a large set of proline substitutions for HIV Env (51). This study provides a promising design candidate for follow-up studies, underscoring the value of the set of previously determined, though somewhat limited, set of E2-bnAb complex structures. Prospective elucidation of the structure of E2 in complex with additional bNAbs, as well as characterization of the E1E2 complex structure, will facilitate future structure-based design studies to engineer and optimize immunogens for an effective HCV vaccine.

## Materials and Methods

### Computational modeling and design

Proline substitution designs to stabilize epitopes were modeled as previously described for design of T cell receptor binding loops (52), using a Ramachandran plot server to assess epitope residue backbone conformations for proline and pre-proline conformational similarities (http://zlab.bu.edu/rama)(53), as well as explicit modeling of energetic effects of proline substitutions using the point mutagenesis mode of Rosetta version 2.3 (54). N-glycan sequon substitutions (NxS, NxT) were modeled using Rosetta (54), followed by modeling of the N-glycan structure using the Glyprot web server (55). Assessment of residue side chain accessible surface areas was performed using NACCESS (56) with default parameters.

### Protein and antibody expression and purification

Expression and purification of recombinant soluble HCV E2 (sE2) and designs was performed as previously described (19). Briefly, the sequence from isolate H77C (GenBank accession number AF011751; residues 384–661) was cloned into the pSecTag2 vector (Invitrogen), transfected with 293fectin into FreeStyle HEK293-F cells (Invitrogen), and purified from culture supernatants by sequential HisTrap Ni^2+^-NTA and Superdex 200 columns (GE Healthcare). For recombinant HCV E1E2 expression, the H77C E1E2 glycoprotein coding region (GenBank accession number AF011751) was synthesized with a modified tPA signal peptide (57) at the N-terminus and cloned into the vector pcDNA3.1+ at the cloning sites of KpnI/NotI (GenScript). Expi293 cells (Thermo Fisher) were used to express the E1E2 glycoprotein complex. In brief, the Expi293 cells were grown in Expi293 medium (ThermoFisher) at 37°C, 125 rpm, 8% CO2 and 80% humidity in Erlenmeyer sterile polycarbonate flasks (VWR). The day before the transfection, 2.0 × 10^6^ viable cells/ml was seeded in a flask and the manufacturer’s protocol (A14524, ThermoFisher) was followed for transfection performance. After 72 hours post-transfection, the cell pellets were harvested by centrifuging cells at 3,000 x g for 5 min and the cell pellet were then stored at −80 °C for further processing. Recombinant E1E2 was extracted from cell membranes using 1% NP-9 and purified via sequential Fractogel EMD TMAE (Millipore), Fractogel EMD SO_3_^−^ (Millipore). HC84.26 immunoaffinity, and Galanthus Nivalis Lectin (GNL, Vector Laboratories) affinity chromatography. Monoclonal antibody HCV1 was provided by Dr. Yang Wang (MassBiologics, University of Massachusetts Medical School), and monoclonal antibodies AR3A, AR4A, and AR5A were provided by Dr. Mansun Law (Scripps Research Institute). All other monoclonal antibodies used in ELISA and binding studies were produced as previously described (24, 25, 58). A clone for mammalian expression of CD81 large extracellular loop (LEL), containing N-terminal tPA signal sequence and C-terminal twin Strep tag, was provided by Joe Grove (University College London). CD81-LEL was expressed through transiently transfection in Expi293F cells (ThermoFisher) and purified from supernatant with a Gravity Flow Strep-Tactin Superflow high capacity column (IBA Lifesciences). Purified CD81-LEL was polished by size exclusion chromatography (SEC) with a Superdex 75 10/300 GL column (GE Healthcare) on an Akta FPLC (GE Healthcare).

### ELISA antigenic characterization and competition assays

Cloning and characterization of E2 mutant antigenicity using ELISA was performed as described previously (28). Mutants were constructed in plasmids carrying the 1a H77C E1E2 coding sequence (GenBank accession number AF009606), as described previously (59). All the mutations were confirmed by DNA sequence analysis (Elim Biopharmaceuticals, Inc., Hayward, CA) for the desired mutations and for absence of unexpected residue changes in the full-length E1E2-encoding sequence. The resulting plasmids were transfected into HEK 293T cells for transient protein expression using the calcium-phosphate method. Individual E2 protein expression was normalized by binding of CBH-17, an HCV E2 HMAb to a linear epitope (60). Data are shown as mean values of two experiments performed in triplicate.

Serum samples at specified dilutions were tested for their ability to block the binding of selected HCV HMAbs-conjugated with biotin in a GNA-captured E1E2 glycoproteins ELISA, as described (24).

### Biolayer interferometry

The interaction of recombinant sE2 glycoproteins with CD81 and HMAbs in was measured using an Octet RED96 instrument and Ni^2+^-NTA biosensors (Pall ForteBio). The biosensors were loaded with 5 μg/mL of purified His_6_-tagged wild-type or mutant sE2 for 600 sec. Association for 300 sec followed by dissociation for 300 sec against a 2-fold concentration dilution series of each antibody was performed. Data analysis was performed using Octet Data Analysis 10.0 software and utilized reference subtraction at 0 nM antibody concentration, alignment to the baseline, interstep correction to the dissociation step, and Savitzky-Golay fitting. Curves were globally fitted based on association and dissociation to obtain *K*_D_ values.

### Differential scanning calorimetry

Thermal melting curves for monomeric E2 proteins were acquired using a MicroCal PEAQ-DSC automated system (Malvern Panalytical). Purified monomeric E2 proteins were dialyzed into PBS prior to analysis and the dialysis buffer was used as the reference in the experiments. Samples were diluted to 10 μM in PBS prior to analysis. Thermal melting was probed at a scan rate of 90 °Ch^−1^ over a temperature range of 25 to 115 °C. All data analyses including estimation of the melting temperature were performed using standard protocols that are included with the PEAQ-DSC software.

### Mass spectrometry

Digestion was performed on 40 μg each of HEK293-derived sE2 glycan sequon substitutions by denaturing using 6 M guanidine HCl, 1 mM EDTA in 0.1 M Tris, pH 7.8, reduced with a final concentration of 20 mM DTT (65 °C for 90 min), and alkylated at a final concentration of 50 mM iodoacetamide (room temperature for 30 min). Samples were then buffer exchanged into 1 M urea in 0.1 M Tris, pH 7.8 for digestion. Sequential digestion was performed using trypsin (1/50 enzyme/protein ratio, w/w) for 18 hours at 37 °C, followed by chymotrypsin (1/20 enzyme:protein, w/w) overnight at room temperature. Samples were then absorbed onto Sep-Pak tC18 columns to remove proteolytic digestion buffer, eluted with 50% acetonitrile/0.1% trifluoroacetic acid (TFA) buffer and concentrated to dryness in a centrifugal vacuum concentrator. The samples were then resuspended in 50 mM Sodium acetate pH 4.5 and incubated with Endo F1, Endo F2, and Endo F3 (QA Bio) at 37 °C for 72 hours to remove complex glycans. LC-UV-MS analyses were performed using an UltiMate 3000 LC system coupled to an LTQ Orbitrap Discovery equipped with a heated electrospray ionization (HESI) source and operated in a top 5 dynamic exclusion mode. A volume of 25 μl (representing 10 μg of digested protein) of sample was loaded via the autosampler onto a C18 peptide column (AdvanceBio Peptide 2.7 um, 2.1 x 150 mm, Agilent part number 653750-902) enclosed in a thermostatted column oven set to 50 °C. Samples were held at 4 °C while queued for injection. The chromatographic gradient was conducted as described previously (Urbanowitz et al). Identification of glycosylated peptides containing the glycan sequon substitution was performed using Byonic software and extracted ion chromatograms used for estimating the relative abundance of the glycosylated peptides in Byologic software (Protein Metrics).

### Animal immunization

CD-1 mice were purchased from Charles River Laboratories. Prior to immunization, sE2 antigens were formulated with polyphosphazene adjuvant. Poly[di(carboxylatophenoxy)phosphazene], PCPP (molecular weight 800,000 Da) (61) was dissolved in PBS (pH 7.4) and mixed with sE2 antigen solution at 1:1 (prime) or 1:5 (w/w) (boost immunization) antigen:adjuvant ratio to provide for 50 mcg PCPP dose per animal. The absence of aggregation in adjuvanted formulations was confirmed by dynamic light scattering (DLS): single peak, z-average hydrodynamic diameter – 60 nm. The formation of sE2 antigen – PCPP complex was proven by asymmetric flow field flow fractionation (AF4) as described previously (62). On scheduled vaccination days, groups of 6 female mice, age 7-9 weeks, were injected via the intraperitoneal (IP) route with a 50 µg sE2 prime (day 0) and boosted with 10 µg sE2 on days 7, 14, 28, and 42. Blood samples were collected prior to each injection with a terminal bleed on day 56. The collected samples were processed for serum by centrifugation and stored at −80°C until analysis was performed.

### Serum peptide and protein ELISA

Domain-specific serum binding was tested using ELISA with C-terminal biotinylated peptides from H77C AS412 (aa 410-425; sequence NIQLINTNGSWHINST) and AS434 (aa 434-446; sequence NTGWLAGLFYQHK), using 2 μg/ml coating concentration. Recombinant sE2 and E1E2 proteins were captured onto GNA-coated microtiter plates. Endpoint titers were calculated by curve fitting in GraphPad Prism software, with endpoint OD defined as four times the highest absorbance value of Day 0 sera.

### HCV pseudoparticle generation

HCV pseudoparticles (HCVpp) were generated as described previously (19), by co-transfection of HEK293T cells with the murine leukemia virus (MLV) Gag-Pol packaging vector, luciferase reporter plasmid, and plasmid expressing HCV E1E2 using Lipofectamine 3000 (Thermo Fisher Scientific). Envelope-free control (empty plasmid) was used as negative control in all experiments. Supernatants containing HCVpp were harvested at 48 h and 72 h post-transfection, and filtered through 0.45 μm pore-sized membranes. Concentrated HCVpp were obtained by ultracentrifugation of 33 ml of filtered supernatants through a 7 ml 20% sucrose cushion using an SW 28 Beckman Coulter rotor at 25,000 rpm for 2.5 hours at 4°C, following a previously reported protocol (26).

### HCVpp serum binding

For measurement of serum binding to HCVpp, 100 µL of 0.45 µm filtered HCVpp isolates were directly coated onto Nunc-immuno MaxiSorp (Thermo Scientific) microwells overnight at 4°C. Microwells were washed three times with 300 µL of 1X PBS, 0.05% Tween 20 in between steps. Wells were blocked with Pierce Protein-Free Blocking buffer (Thermo Scientific) for 1 hour. Serum sample dilutions made in blocking buffer were added to the microwells and incubated for 1 hour at room temperature. Abs were detected with secondary HRP conjugated goat anti-mouse IgG H&L (Abcam, ab97023) and developed with TMB substrate solution (Bio-Rad). The reaction was stopped with 2N sulfuric acid. A Molecular Devices M3 plate reader was used to measure absorbance at 450 nm. Endpoint titers were calculated by curve fitting in GraphPad Prism software, with endpoint OD defined as four times the highest absorbance value of Day 0 sera.

### HCVpp neutralization assays

For infectivity and neutralization testing of HCVpp, 1.5 × 10^4^ Huh7 cells per well were plated in 96-well tissue culture plates (Corning) and incubated overnight at 37 °C. The following day, HCVpp were mixed with appropriate amounts of antibody and then incubated for 1 h at 37 °C before adding them to Huh7 cells. After 72 h at 37 °C, either 100 μl Bright-Glo (Promega) was added to each well and incubated for 2 min or cells were lysed with Cell lysis buffer (Promega E1500) and placed on a rocker for 15 min. Luciferase activity was then measured in relative light units (RLUs) using either a SpectraMax M3 microplate reader (Molecular Devices) with SoftMax Pro6 software (Bright-Glo protocol) or wells were individually injected with 50 μL luciferase substrate and read using a FLUOstar Omega plate reader (BMG Labtech) with MARS software. Infection by HCVpp was measured in the presence of anti-E2 MAbs, tested animal sera, pre-immune animal sera, and non-specific IgG at the same dilution. Each sample was tested in duplicate or triplicate. Neutralizing activities were reported as 50% inhibitory dilution (ID_50_) values and were calculated by nonlinear curve fitting (GraphPad Prism), using lower and upper bounds (0% and 100% inhibition) as constraints to assist curve fitting.

### Statistical comparisons

P-values between group endpoint titers and group ID_50_ values were calculated using Kruskal-Wallis one-way analysis of variance (ANOVA), with Dunn’s multiple comparison test, in Graphpad Prism software.

## Acknowledgements

We thank Joe Grove (University College London) for kindly providing the CD81-LEL expression plasmid. We also thank Verna Frasca (Malvern Panalytical) for performing and analyzing the DSC experiments, and Sneha Rangarajan (University of Maryland IBBR) for useful discussions regarding the antigenic domain D structure. This work was supported in part by National Institute of Allergy and Infectious Diseases/NIH grants R21-AI126582 (BGP, RAM, SKHF), R01-AI132213 (BGP, AKA, RAM, TRF, SKHF), and U19-AI123862 (SKHF).

---

**Figure S1.**
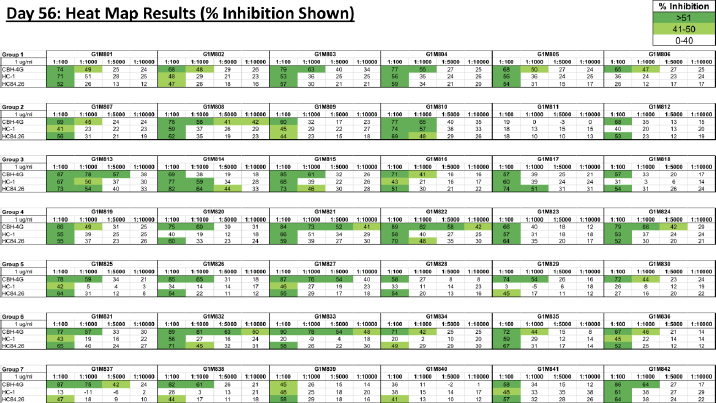
Serum binding competition with monoclonal antibodies. Serum binding competition with monoclonal antibodies CBH-4G, HC-1, and HC84.26 was tested at the serum dilutions shown, using ELISA.

**Figure S2.**
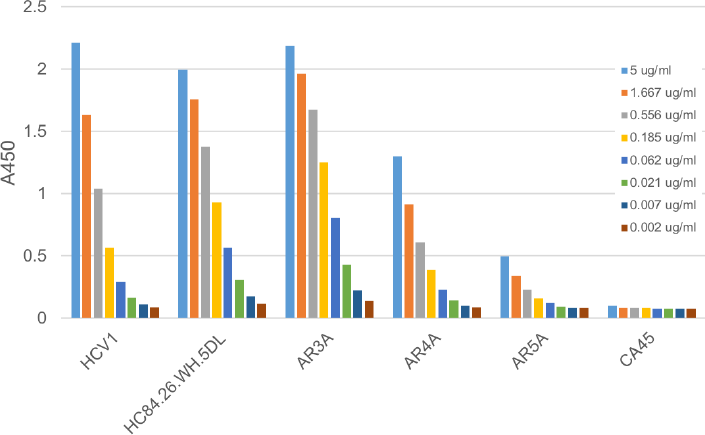
Binding of purified HCV pseudoparticles (HCVpps), pseudotyped with H77C E1E2, to monoclonal antibodies. Binding measurements were performed using ELISA with antibodies targeting E2 (HCV1, HC84.26.WH.5DL, AR3A), E1E2 (AR4A, AR5A) and a negative control antibody (CA45).

